# Cardiac function and ECM morphology are altered with high fat diets in *Drosophila*

**DOI:** 10.1101/2023.08.08.552539

**Authors:** Rachel M. Andrews, Saumya Naik, Katie Pelletier, J. Roger Jacobs

## Abstract

Cardiovascular disease is characterized by aberrant and excessive extracellular matrix (ECM) remodelling, termed fibrosis. Fibrotic remodelling is typically triggered by inflammation, which occurs systemically in obesity. Despite the contribution of fibrosis to adverse clinical outcomes and disease progression, there are no available treatments for this condition. Developing therapeutics for chronic conditions requires an understanding of *in vivo* ECM regulation, and how the ECM responds to a systemic challenge. We have therefore developed a *Drosophila* model for obesity via chronic high fat diet feeding and evaluated the response of the cardiac ECM to this metabolic challenge. We found that this model displays a striking disorganization of the cardiac ECM, with corresponding deficits in heart function. Our study shows that different genotypes tolerate varying levels of high fat diets, and that some genotypes may require a different percentage of fat supplementation for achieving an optimal obesity phenotype.

## Introduction

Cardiovascular disease (CVD) is a leading cause of death world-wide. In recent years, there has been an increase in age-adjusted mortality resulting from CVD (Sidney et al. 2022). Obesity is one of the main risk factors for the development of CVD, and an increased incidence of obesity has led to a corresponding increase in CVD rates (Poirier et al. 2006). One aspect of CVD that is often overlooked is the contribution of the extracellular matrix (ECM). The ECM is a protein scaffold that surrounds tissues within the body and acts to support their function by modulation of tension, distribution of forces through the tissue, sequestration of growth factors, and, of importance in the heart, mediation of electrical conduction (Travers et al. 2016; Bonnans, Chou, and Werb 2014; Cox and Erler 2011). The importance of the ECM is demonstrated by its intimate link to a wide variety of disease states, including cancer and CVD.

The ECM is composed of two main compartments, the interstitial matrix and the basement membrane. The interstitial matrix is composed primarily of fibrillar Collagen and forms a support scaffold, while the basement membrane is found close to the cell surface and acts as a barrier to the surroundings (Bonnans, Chou, and Werb 2014; Walker and Spinale 1999). The ECM is not a static structure, and undergoes constant turnover, called remodelling. This process is a fine-tuned balance of matrix synthesis and deposition, as well as matrix breakdown (Cox and Erler 2011; Hughes and Jacobs 2017). The matrix metalloproteinases (MMPs) are mainly responsible for ECM breakdown, and their level of activity in the tissue is regulated by their inhibitors, the tissue inhibitors of metalloproteinases (TIMPs) (Hughes et al. 2020). The ratio of matrix deposition to breakdown is an important contributor to the biophysical properties of a tissue (Cox and Erler 2011; Mouw, Ou, and Weaver 2014).

The basement membrane is made up of highly conserved core components. It consists of a Laminin sheet that is anchored to the cell surface by Integrins. A second sheet of Collagen-IV is then anchored to the Laminin sheet, primarily through cross-linking proteins like Nidogen (Hughes and Jacobs 2017; Howard et al. 2019). The basement membrane is an ancient structure that is found in all metazoans and is thought to have facilitated the development of multicellularity (Fidler et al. 2017). The majority of metazoans also possess the fibrillar collagens that make up the interstitial matrix. One notable exception is the fruit fly, *Drosophila melanogaster*. *Drosophila* lacks the common fibrillar collagens, making it an intriguing model for the study of basement membrane dynamics.

Due to its critical involvement in supporting and maintaining tissue function, ECM homeostasis is tightly controlled (Bonnans, Chou, and Werb 2014). One of the results of ECM dysregulation is fibrotic remodelling, or fibrosis. Fibrosis refers to increased deposition of matrix components, as well as increased levels of crosslinking between these proteins (Travers et al. 2016; Meschiari et al. 2017). In a healthy ECM there is a balance of protein breakdown and deposition that acts to maintain tissue characteristics. With fibrosis, this balance is disrupted in favour of increased protein deposition. Increased levels of crosslinking insolubilize the matrix and make it more resistant to degradation, further disrupting the normal balance of remodelling.

In the heart specifically, fibrosis can have catastrophic consequences due to the replacement of highly specialized, contractile cardiomyocytes with non-contractile ECM proteins. This compromises the ability of the heart to contract effectively, can disrupt the connections between cells that are crucial for conduction of nerve impulses, and leads to maladaptive cardiac remodelling that can progress to heart failure (Travers et al. 2016). Cardiac fibrosis is initially adaptive, with activated fibroblasts laying down additional ECM proteins in order to replace necrotic cells and prevent cardiac rupture (Travers et al. 2016). However, as these cells persist in the tissue, fibrosis becomes progressive and heart function is further compromised (Jourdan-LeSaux, Zhang, and Lindsey 2010).

Despite being a prevalent component of many diseases, fibrosis has no available treatments (Pehrsson et al. 2021). Essentially every case of CVD exhibits some level of fibrotic remodelling (Travers et al. 2016; El Hajj et al. 2018). Fibrotic remodelling of the heart is also known to occur in the context of obesity, one of the leading comorbidities of heart disease (Cavalera, Wang, and Frangogiannis 2014). This necessitates a deeper understanding of how the cardiac ECM responds to obesity specifically, which could allow for more targeted treatment of disease symptoms.

In order to identify the specific effects of obesity on the cardiac ECM, we have employed *Drosophila melanogaster. Drosophila* is a powerful tool for performing this research due to its lack of genetic redundancy, simple heart tube that is not required to support life, and a similar ECM to mammals, including the presence of highly conserved basement membrane proteins (Hughes and Jacobs 2017; Pastor-Pareja 2020; Rotstein and Paululat 2016). The *Drosophila* heart, or dorsal vessel, is a linear, tube-like structure that follows the same developmental pathways as the early mammalian heart. The *Drosophila* heart has no stem cells so has no replacement of cells after damage (Hughes and Jacobs 2017). While the mammalian heart does have some stem cells they do not contribute meaningfully to cardiomyocyte replacement (van Berlo et al. 2014). Following injury the mammalian heart relies on repair rather than replacement of cells (Vujic, Natarajan, and Lee 2020). Studying repair mechanisms in the *Drosophila* heart may therefore provide insights into how mammalian hearts function following injury.

The *Drosophila* cardiac ECM is composed of the same core components as other basement membranes. It contains both Laminin and Collagen-IV, as well as the heart-specific collagen Pericardin. Pericardin is a Collagen-IV like protein that organizes similarly to fibrillar collagens in mammals (Chartier et al. 2002; Sessions et al. 2017). Thus, the *Drosophila* cardiac ECM is composed of two Collagen matrices, one of Collagen-IV and one of the more fibrillar appearing Pericardin. Pericardin has previously been shown to be critical for the maintenance of the *Drosophila* heart (Chartier et al. 2002; Drechsler et al. 2013).

Diet has been used previously to induce obesity-like phenotypes in *Drosophila,* including methods that supplement a standard *Drosophila* diet with coconut oil as a source of fat and excess calories (Birse et al. 2010; Diop, Birse, and Bodmer 2017). The present study utilized a dosage series of high fat diets (HFDs) to determine what affect dietary supplementation had on the *Drosophila* heart. Previous studies in adult *Drosophila* have revealed cardiac dysfunction, increased triglyceride levels, and altered metabolism as a result of short-term HFD feeding (Guida et al. 2019; Birse et al. 2010; Diop, Birse, and Bodmer 2017). The present study aimed to expand on these results by feeding larval *Drosophila* a HFD from hatching. Larvae were allowed to feed until late third instar in order to maximize the duration of dietary treatment, producing a chronic high fat diet model. In *Drosophila* growth and maturation are distinct stages, with growth occurring exclusively during larval stages (Rewitz, Yamanaka, and O’Connor 2013). By performing these experiments during a growth phase of the life cycle we are able to administer the HFD chronically, circumvent egg-laying and other adult behaviours, and maximally stress the heart as it must grow and adapt to HFD conditions. Utilizing larval stages also eliminates the confound of aging, which naturally leads to cardiac ECM accumulation (Sessions et al. 2017; Hinderer and Schenke-Layland 2019). Additionally, studies conducted on adult *Drosophila* heart function typically examine only females due to their larger body size (Birse et al. 2010; Guida et al. 2019; Walls et al. 2020). Focusing on the larval life stage allows for analysis of both female and male larvae as size dimorphisms are more limited during the growth stage. This allows us to examine any sex-specific effects of HFD treatments more easily than in adult *Drosophila*.

Additionally, a high sucrose diet was employed in this study to determine if the type of nutrient providing the excess calories affected the heart specifically. High sucrose diets have been shown to induce a diabetes-like phenotype in *Drosophila*, exhibiting insulin resistance, cardiac arrhythmias, and accumulation of Pericardin (Na et al. 2013). This dietary treatment allows for a distinction to be made between excess caloric intake or inducing an obesity phenotype as the cause of cardiac dysfunction.

Here, we describe the effects of chronic HFD feeding on the larval *Drosophila* cardiac ECM. We observed that HFD results in several hallmarks of obesity, and causes changes to matrix organization of both Pericardin and Collagen-IV. Pericardin organization was severely perturbed, with the protein network revealing an anterior-posterior fibre alignment phenotype that was rarely observed in controls. The Collagen-IV matrix demonstrated a clumping phenotype, with a dose-dependent level of clumping within the matrix. We also observed functional impairment of the heart, namely an inability to contract fully at systole. This could be due to the rearrangement of the cardiac ECM, and altered tension through the dorsal vessel. Overall, our results suggest that HFD feeding in *Drosophila* larvae induces an obesity-like phenotype that affects the organization of the cardiac ECM and leads to cardiovascular impairment.

## Results

### Viability of larvae on high fat and high sucrose diets

To determine the dilution series of HFD treatments larvae were reared from hatching on food containing 10%-50% coconut oil. 50% had no larvae survive to late L3 (data not shown) so 10%-40% dosages were used for all experiments. 1M sucrose was selected based on previous studies (Palanker Musselman et al. 2011) and is calorically comparable to 20% HFD supplementation. Larvae did not survive a 5M diet which would have been equivalent in calories to the 40% HFD treatment (data not shown).

Health of larvae was assessed by measuring larval mass, triglyceride levels, and lipid droplet diameter. All assays separated female and male larvae in order to isolate possible sex specific effects. Assays were also conducted on both *y^1^w^1118^* and *y^1^w^1118^ ; vkg-GFP* (hereafter *vkgGFP*) so both Collagens in the cardiac ECM could be examined. *VkgGFP* is a protein trap that labels the *viking* subunit of Collagen-IV with a GFP tag, so endogenous Collagen-IV is fluorescently tagged (Buszczak et al. 2007). Larval mass was mildly elevated in 10% and 20% female *y^1^w^1118^* high fat diet treatment groups, and 10%-30% in males (elevation between 13.8-14.8% in females, and 15.1-20.6% in males) (**Figure 1A**). A modest increase or unchanged body size was not unexpected as the critical weight checkpoint triggers the transition from larval development to pupation (Zeng et al. 2020). An increase in weight could therefore trigger early pupation rather than continued increase in larval size. The high sucrose diet resulted in larvae that were significantly smaller controls and the calorically equivalent HFD treatment (females 43.6% mass of controls, males 54%) (**Figure 1A**). This is consistent with previous studies that demonstrate high sucrose inducing diabetes-like phenotypes as a result of impaired insulin signalling (Na et al. 2013).

**Figure 1:**
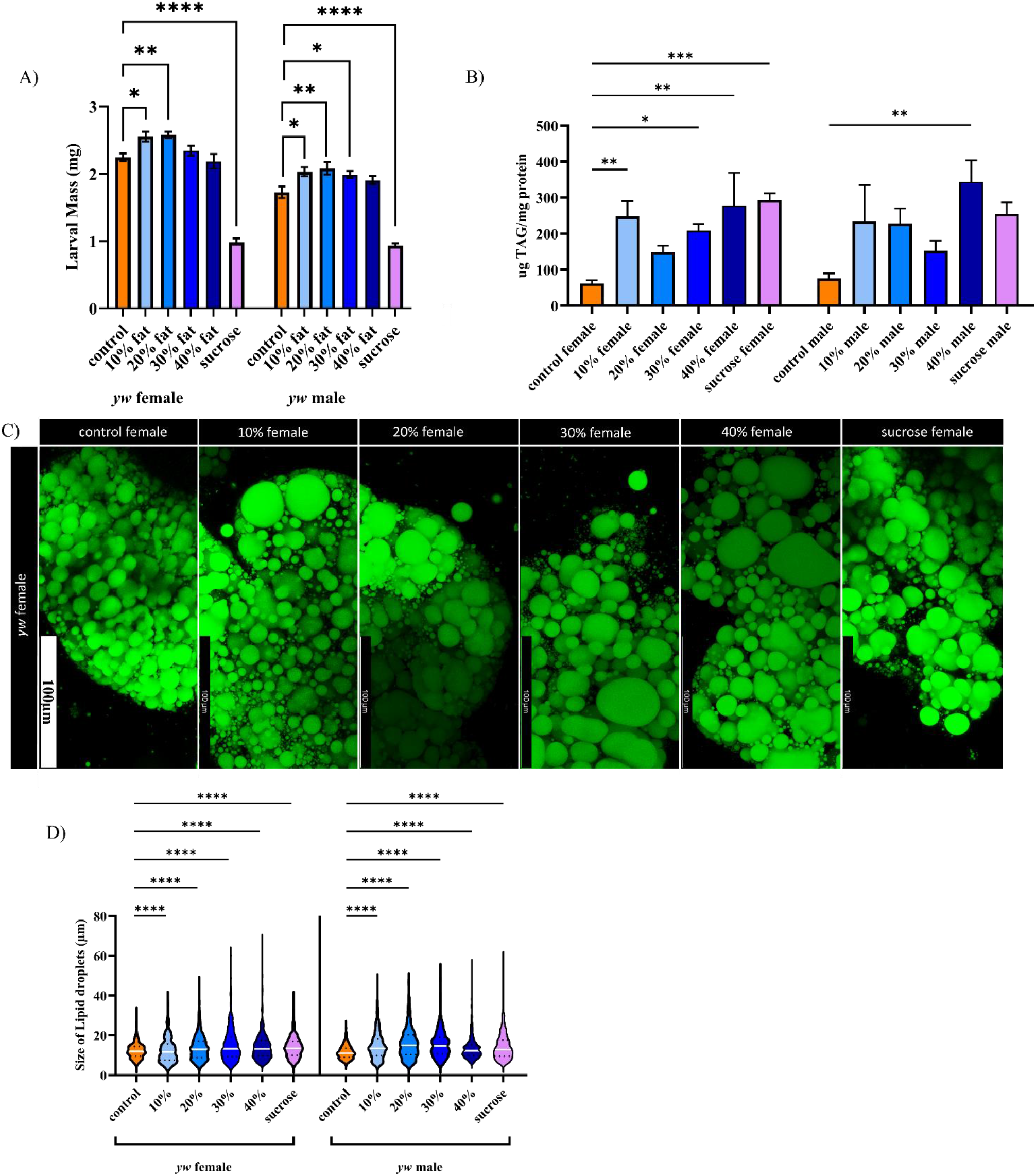
Larvae fed a high fat diet show a dose dependent increase in markers of obesity. Larval mass is not lowered by high fat diet feeding, in contrast to high sucrose (A). HFD results in a dose dependent increase in triglyceride levels in *y^1^w^1118^* larvae, especially females (B). Lipid droplets were visualized with BODIPY 493/503 (C) and reveal that both HFD feeding and a high sucrose diet result in a dose-dependent increase in lipid droplet size (D). Error bars in A and B are SEM. White lines in D represent the median, dotted lines represent quartiles. *=p<0.05, **=p<0.01, ***=p<0.001, ****=p<0.0001

In *vkgGFP* larvae 20% and 30% HFD females were smaller than the control genotype (control=2.267mg+/-0.0335mg, 20%=1.931mg+/-0.0468, 30%=1.959mg+/-0.0895). This suggests that *vkgGFP* does not tolerate the HFD treatment as well as the *y^1^w^1118^* genotype (**Figure S1**).

Triglyceride levels were measured and *y^1^w^1118^* larvae exhibited a dose dependent increase in triglyceride levels compared to controls (**Figure 1B**). The overall trend was similar for both female and male larvae but female larvae exhibited a more significant change. *vkgGFP* larvae did not demonstrate an elevation in triglyceride level but controls of this genotype had triglyceride levels over 4 times higher than *y^1^w^1118^* controls (**Figure S2**). Oregon R triglyceride levels were intermediate between *y^1^w^1118^* and *vkgGFP,* suggesting that triglyceride level can vary markedly with genotype (**Figure S2**). The higher baseline triglyceride level in *vkgGFP* individuals may contribute to their reduced ability to tolerate HFD feeding when compared to *y^1^w^1118^.* This may also indicate that there is a limit to the level of triglycerides that larvae are able to process effectively, and *vkgGFP* does not have a significant increase in triglyceride levels with HFD feeding because they are already close to this limit at baseline.

Lipid droplet diameter in *y^1^w^1118^* individuals showed a dose dependent effect with HFD feeding (**Figure 1 C).** All HFD had increased lipid droplet diameter (**Figure 1D**). The high sucrose diet also exhibited significantly increased droplet diameter.

Overall this shows that high fat diets are able to induce obesity-like phenotypes, while high sucrose diets have some characteristics of obesity but are significantly smaller than controls and HFD treatments. Previous studies have shown that this is due to changes in insulin signalling that induces a diabetes-like phenotype in these individuals (Palanker Musselman et al. 2011).

### Pericardin fibre organization

Having established that HFD feeding results in several hallmarks of obesity, we investigated the effect of HFD feeding on the cardiac ECM. The fibrous, heart specific collagen Pericardin showed marked changes in its organization with all dietary treatments. A normal Pericardin matrix is an organized meshwork that forms a honey-comb like pattern and extends away from the heart tube (**Figure 2A, A’**). With dietary treatments, the matrix takes on an anterior-posterior linearity phenotype (**Figure 2B-F’**) that is rarely observed in controls. Both *y^1^w^1118^*and *vkgGFP* larvae demonstrate this phenotype with all dietary treatments (**Figure 3A**). Interestingly, in *vkgGFP* specifically, there is a slight improvement in the percentage of the population that is affected at 40% feeding (**Figure 4F**). This was likely due to a survivor bias in this group. Percent survival of the *y^1^w^1118^*genotype was relatively unaffected at 30% and 40% HFD but was significantly reduced in *vkgGFP,* especially at 40%. *vkgGFP* appears to be less able to tolerate the highest HFD treatments than *y^1^w^1118^*larvae.

**Figure 2:**
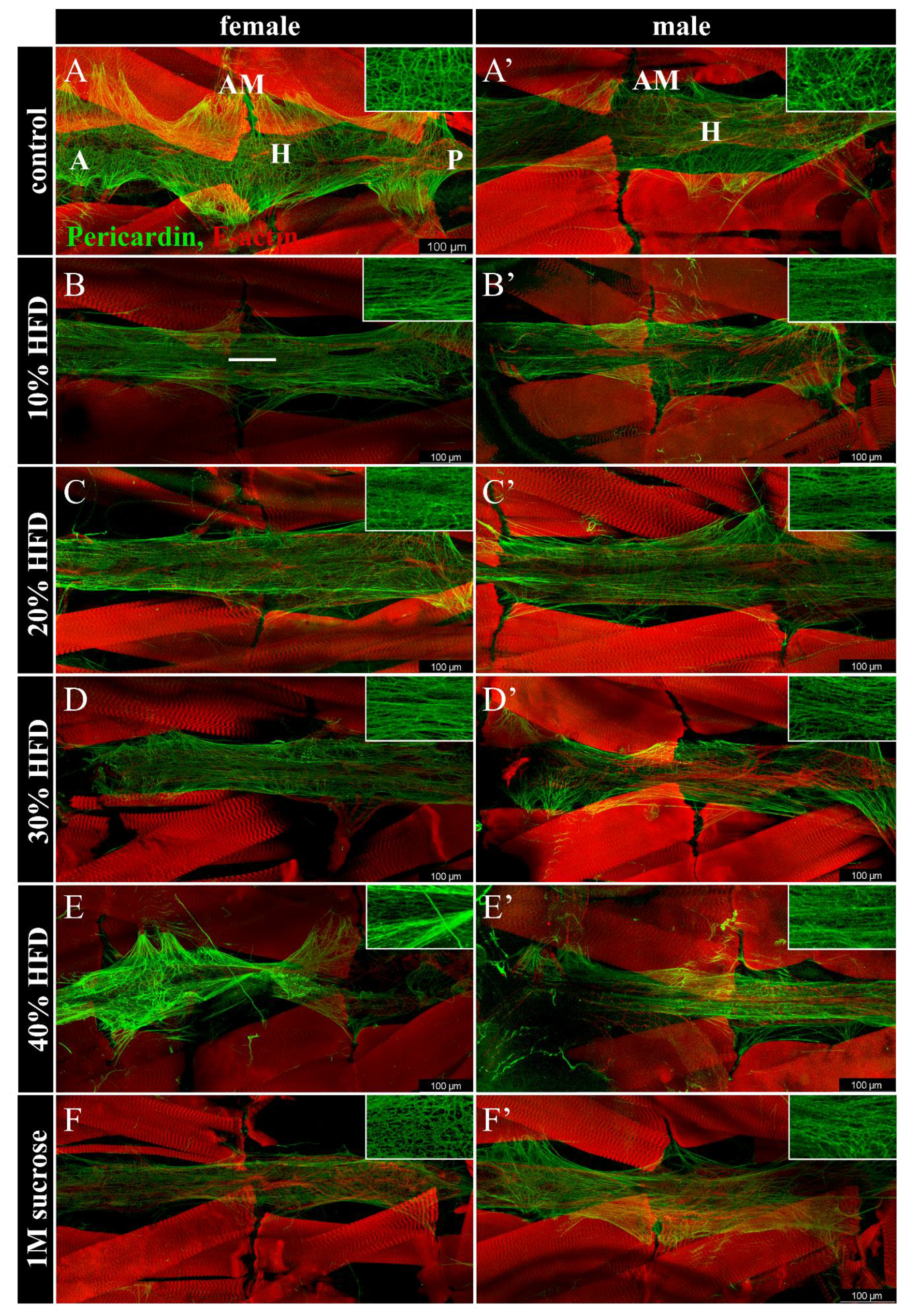
Pericardin fibre organization is perturbed in *y^1^w^1118^* dietary treatments. Controls demonstrate a normal, organized meshwork (A-A’) while dietary treatments show a change in matrix organization, with matrix fibres becoming oriented anterior-posterior (B-F’). The cardiac ECM is visualized by immunolabelling Pericardin (green) and F-actin (red). The F-actin label in the background is the body wall muscles. In panel A, H labels the heart tube, AM labels alary muscles, A is anterior, P is posterior. All images are oriented anterior to the left. White line in panel B follows direction of anterior-posterior orientation of Pericardin fibres.

**Figure 3:**
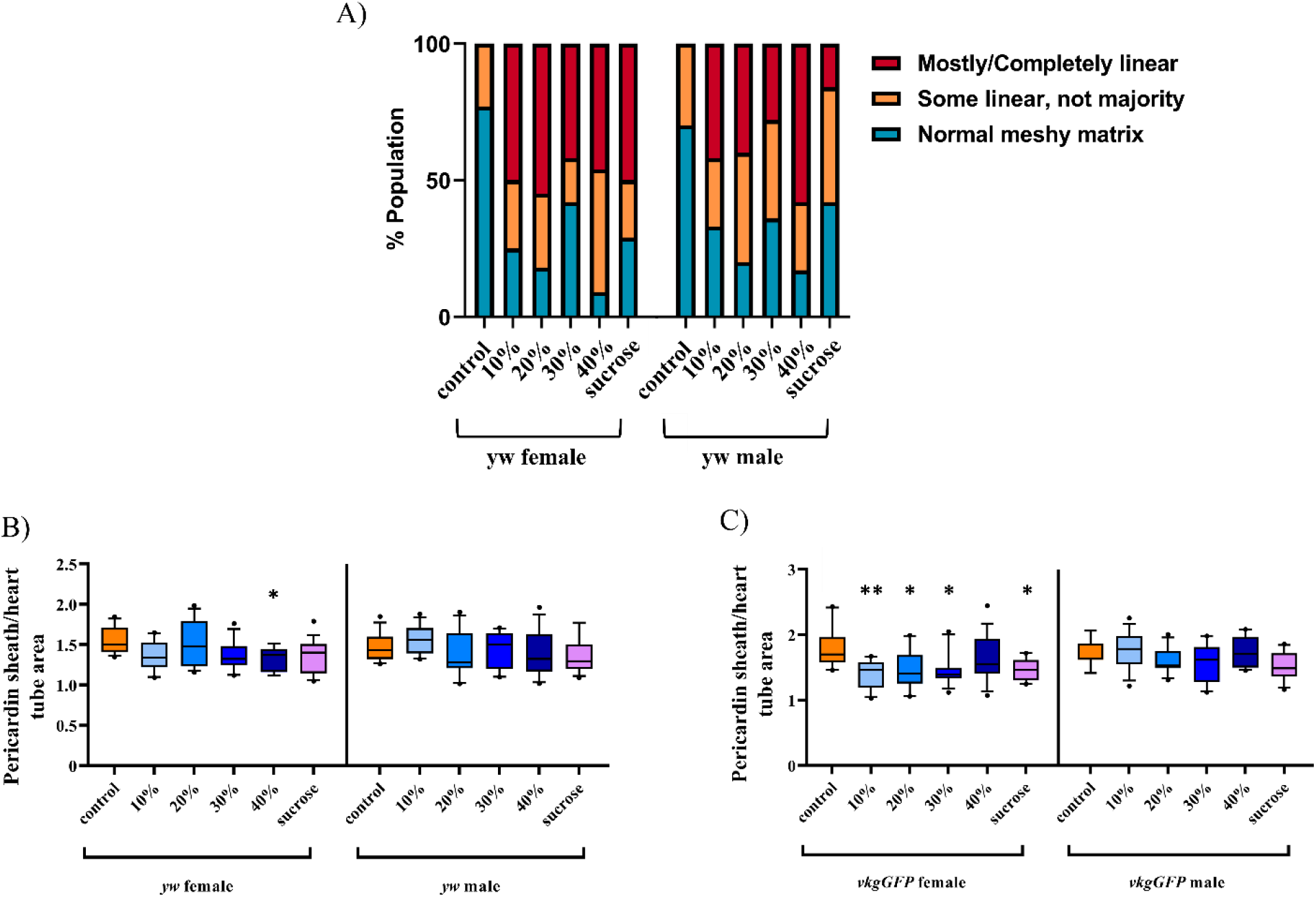
Matrix organization shows significant rearrangement in dietary treatments. Dietary treatments demonstrate a linearity phenotype that is more common and more severe than controls (A). In females, there is a downward trend in the size of the Pericardin matrix relative to the heart tube (B), not statistically significant, and (C) but not in males. *=p<0.05, **=p<0.01

**Figure 4:**
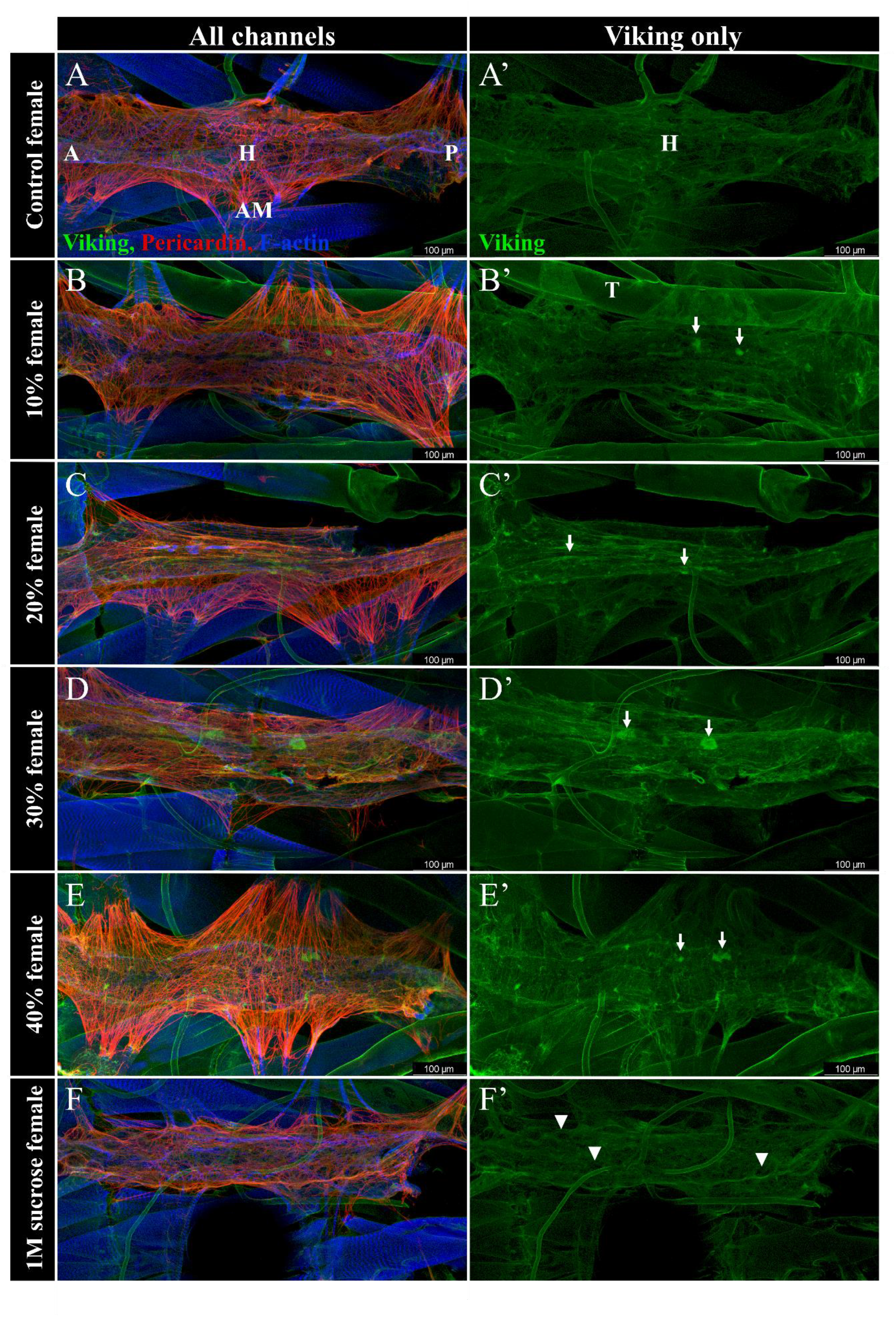
Collagen-IV distribution is abnormal in *vkgGFP* dietary treatments. In control individuals the Collagen-IV matrix shows a regular, sheet-like distribution across the surface of the heart (A’). HFD treated larvae experience elevated levels of clumping within the Collagen-IV matrix, indicated by white arrows, that is not found in the high sucrose diet treatment (B’,C’, D’, E’). Holes are observed in the high sucrose Collagen-IV matrix instead, indicated by white arrowhead (F’). The cardiac ECM is visualized by endogenous *vkgGFP* fluorescence (green), and immunolabelling Pericardin (red) and F-actin (blue). In panel A, H labels the heart tube, AM labels alary muscles, A is anterior, P is posterior. *vkgGFP* is also found in the trachea, indicated by T in panel B’. All images are oriented anterior to the left.

The Pericardin matrix also appears to not extend away from the heart tube with high fat diet feeding (**Figure 2B-F’).** This could be due to altered tension due to changes in matrix organization. The ratio of the area of the Pericardin matrix to the area of the heart tube revealed a clear dose-dependent downward trend in the female *vkgGFP* larvae (**Figure 3C**). A similar trend was observed in the female *y^1^w^1118^*larvae but was not statistically significant (**Figure 3B**). However, this trend was not observed in the males of either genotype, suggesting the effects of the high fat diet may cause different defects in matrix organization between female and male larvae.

### Collagen-IV distribution

The Collagen-IV matrix was visualized using the *vkgGFP* line. The distribution of this matrix is normally sheet-like, covering the entire surface of the heart (**Figure 4A’**). The Collagen-IV matrix showed no changes in the high sucrose diet but showed a clumping phenotype in the high fat diet treatments (**Figure 4B’, C’, D’, E’**). The area of the Collagen-IV matrix occupied by clumps was quantified and revealed no significant difference between the controls and the high sucrose diet (**Figure 5A**). The 20% HFD had significantly elevated levels of clumping in the Collagen-IV matrix in contrast to the calorically equivalent high sucrose diet, with 2.29% more clumping in the HFD treatment (95% CI: −3.74, −0.834, p<0.001). The high sucrose diet and the controls were not significantly different, with the high sucrose diet having 1.32% more clumping than controls (95% CI: −2.79, 1.47, p=0.08). Given that the sucrose diet did not have a significant difference in clumping compared to the controls it was excluded from the HFD comparison. The HFD treatments demonstrated a significant dose-dependent effect on clumping phenotype (DF=2, F=20, p<0.001) (**Figure 5B**). While it was found that while diet has an effect on the clumping phenotype, there was no significant sex effect (DF=1, F=1.9891, p=0.164).

**Figure 5:**
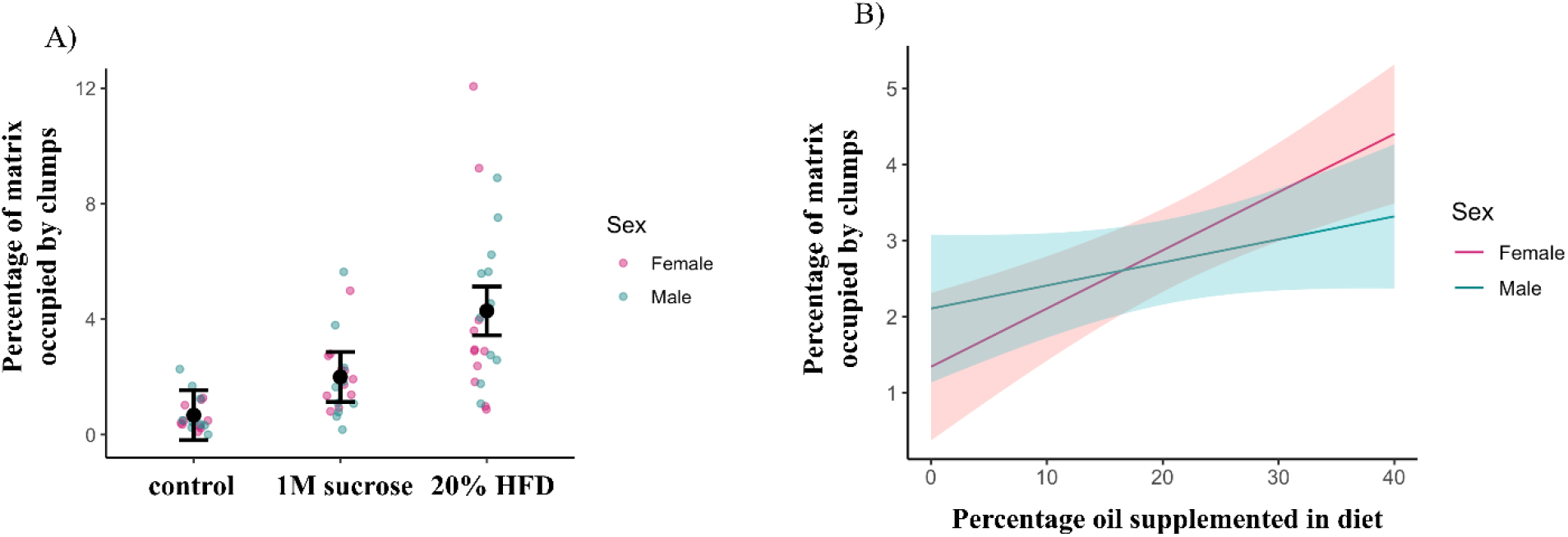
Collagen-IV shows a dose-dependent relationship with clumping defect. High sucrose diet is compared to 20% HFD as they are calorically equivalent. High sucrose diet does not have increased Collagen-IV clumping compared to controls while the calorically comparable HFD does (A). HFD feeding causes a dose-dependent increase in the amount of clumping within the matrix (B). This trend is more pronounced in female larvae. Error bars in A represent 95% confidence intervals. Graph in B is a model estimate with 95% confidence intervals.

A different fibre organization defect was observed in the high sucrose dietary treatment. This group was 1.32% (95% CI: 2.79, 0.15, p=0.08) more likely to have holes or gaps within the normally sheet-like Collagen-IV matrix as compared to the control treatment. This phenotype is not as strong as the clumping phenotype observed in HFD groups. No other treatment group exhibited this phenotype (**Figure S3**). This suggests that the defects in Collagen-IV deposition and remodelling may vary with treatment but this could also be due to reduced growth and altered physiology in the high sucrose group, as major signalling pathways like insulin are known to be affected with this treatment.

### Live imaging by optical coherence tomography

If the cardiac ECM of larvae raised on a HFD reveal perturbations in the organization of both cardiac Collagens how then might this affect cardiac physiology? We performed live imaging to determine if these ECM perturbations have functional consequences using optical coherence tomography (OCT). Movies of beating hearts (see supplemental movies) were used to measure the cross-sectional area of the lumen at both diastole and systole (**Figure 6A**) and it was found that the high sucrose diet generates larvae with much smaller hearts than the controls (**Figure 6C, D**). This is likely due to the smaller body size of these individuals compared to their control counterparts. The high fat diet treatments on the other hand had similar diastolic areas across all treatment groups but showed a dose-dependent increase in systolic area (**Figure 6E**). This indicates an inability of the heart to contract fully with increasing dietary fat, suggesting these larvae have impaired heart function similar to mammalian cases of CVD. However, heart rate was unaffected and arrhythmicity index was only elevated in *y^1^w^1118^* high sucrose diet males **(Figure S4).** Live imaging also revealed that the hearts of HFD treated larvae were more likely to be abnormally shaped (**Figure 6B**). The majority of control hearts were close to round or oval and contracted evenly on all sides. In the high fat diet treated individuals there were irregularly shaped hearts (**Figure 6B**), as well as contraction that mainly occurred in one plane instead of uniformly around the circumference of the heart.

**Figure 6:**
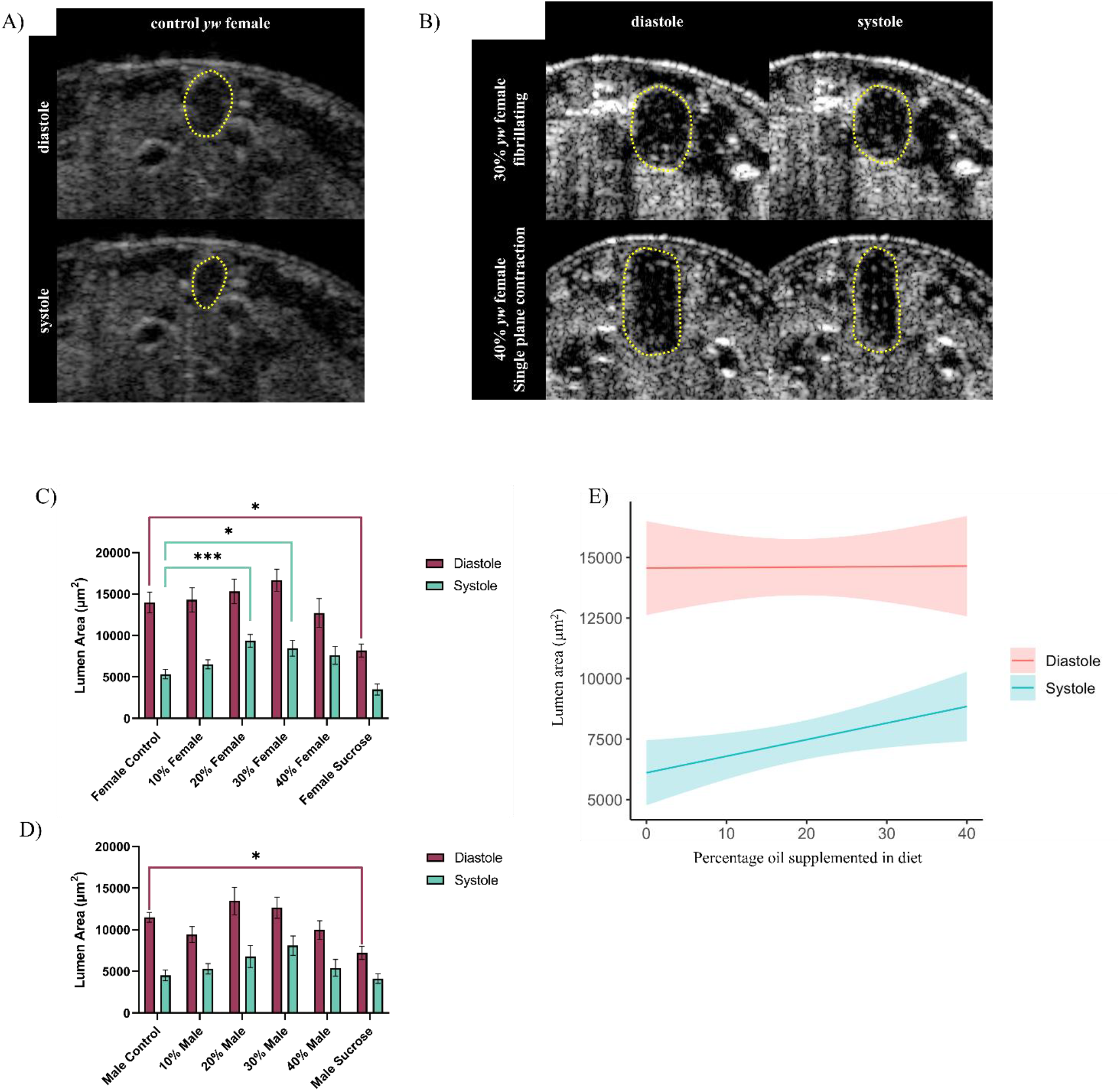
Optical coherence tomography reveals impaired contraction with HFD treatments. OCT can be used to visualize the heart beating in cross-section, revealing the area inside the lumen at both diastole and systole (A). Control hearts show round or oval hearts that contract evenly along the perimeter, while HFD treatments cause abnormal heart shape and can also cause an inability to contract evenly on all sides (B). Lumen cross-sectional area in *y^1^w^1118^* females shows a change in diastole only in high sucrose individuals (C), but does show increases in systolic volume at higher concentrations of HFD. A similar trend is observed in males (D). This demonstrates a dose-dependent impairment of heart contractions (E). Error bars in B and C are SEM. Graph in E is a model estimate with 95% confidence intervals. *=p<0.05, **=p<0.01, ***=p<0.001, ****=p<0.0001

## Discussion

A chronic high fat diet in growing *Drosophila* larvae generates dose-dependent effects on overall organismal health as well as on the form and function of the heart. HFD individuals have clear signs of obesity and CVD, while the high sucrose diet shares some markers of obesity as well as other metabolic disorders. Previous studies in adult *Drosophila* have found that HFD feeding causes increased fat storage, including ectopic deposition of triglycerides, as well as cardiac dysfunction, including decreased diastolic and systolic diameters, reduced fractional shortening (a proxy for stroke volume), and reduced heart period (Hardy et al. 2015; Guida et al. 2019). These health effects point towards the development of lipotoxic cardiomyopathy. Here, we find that larval *Drosophila* experience similar effects, and that the observed cardiac dysfunction may be as a result of altered ECM dynamics.

Genotype was found to affect the magnitude of HFD feeding. *y^1^w^1118^*individuals tolerated the HFD more readily than *vkgGFP* individuals. This may be due to the elevated triglyceride levels found in *vkgGFP* control larvae. These larvae have comparable triglyceride levels to HFD fed *y^1^w^1118^* individuals, suggesting that they are close to a maximum level of tolerance of triglycerides. The increase due to HFD feeding may have caused the reduced viability seen in *vkgGFP* in contrast to *y^1^w^1118^* on HFD treatments. Oregon R triglyceride levels are intermediate to *y^1^w^1118^* and *vkgGFP,* suggesting that triglyceride levels on ordinary lab diets are a variable trait.

HFD feeding affected cardiac ECM organization, with both the Pericardin and Collagen-IV matrices developing defects. Collagen-IV phenotypes for HFD and high sucrose diets were different, with HFD treatments exhibiting a clumping phenotype while high sucrose diets possessed gaps in the matrix. The Collagen-IV clumping phenotype in the HFD feeding treatments suggest that matrix deposition may be elevated. This could be due to increased expression of Collagen-IV or to reduced breakdown (or turnover) of the matrix. Matrix breakdown is performed by the matrix metalloproteinases (MMPs) and altered levels or activity of these proteins have been shown to promote fibrotic remodelling. Altered MMP expression profiles have been demonstrated in a variety of fibrotic diseases (Garrett et al. 2019; Tian, Luo, and Liu 2022). We described previously how depletion of MMP2 by overexpression of its inhibitor TIMP during larval growth causes accumulation of Collagen-IV, suggesting MMP2 expression is required for the maintenance of a healthy ECM (Hughes et al. 2020). Reduced MMP2 activity, either as a result of altered gene expression or due to elevated expression of TIMP, presents an intriguing possibility for the clumping observed the Collagen-IV matrix in these HFD treatment groups.

Organization of the Pericardin network was severely affected by HFD feeding. Instead of forming a honey-comb like meshwork as in controls, HFD feeding and the high sucrose diet were found to induce an anterior-posterior fibre linearity. This linearity phenotype was observed in all HFD treatments. Linearity is rarely seen in controls, suggesting that the ability to organize the matrix appropriately is affected by HFD feeding. This could be due to increased inflammatory responses due to HFD feeding. Excess triglycerides are known to cause an inflammatory response which can lead to remodelling of fibrillar Collagens in cases of CVD in mammalian systems (Cavalera, Wang, and Frangogiannis 2014; Nishida and Otsu 2017). The excess triglycerides measured in HFD fed larvae may be causing a similar inflammatory response that is affecting matrix deposition and turnover, which could in turn lead to the observed reorganization of the Pericardin matrix reported here. ECM regulation is known to be affected in disease states, with elevated levels of crosslinking enzymes, altered post-secretion activation of MMPs and TIMPs, and alterations to proteins important for appropriate Collagen deposition like its chaperone, SPARC (Hartley et al. 2016; Hughes and Jacobs 2017). Our previous study found that MMP activity was critical for the appropriate organization of the Collagen-IV matrix (Hughes et al. 2020). It is therefore feasible that Pericardin organization also requires specific post-translational modifications or enzymatic activity in order to form a normally organized network. If levels of ECM regulators like MMP and SPARC are affected by chronic inflammation, it could affect the overall organization of the Pericardin matrix, generating a matrix with less structural integrity that is prone to collapsing in on itself and appearing linear as observed here.

The typical arrangement of the ECM has Pericardin fibres extending in many directions in order to transmit tension evenly across the heart. This allows for the heart to open evenly in all directions and helps to maintain a uniform shape as the heart contracts and relaxes. Increased linearity of fibres in HFD treated larvae also correlates with defects in the ability of the heart to contract evenly around its perimeter. Both effects suggest that HFD feeding results in an uneven distribution of tension around the heart by affecting the organization of the ECM. Additionally, live imaging by OCT revealed a dose-dependent inability of the heart to contract fully at systole. Diastolic area is unaffected, indicating this defect is in the ability of the heart to contract, not relax. A common finding in CVD in humans is an inability of the heart to contract effectively, including in cases of obesity-related CVD (Alpert, Omran, and Bostick 2016). The functional defects observed in *Drosophila* larvae with an obesity-like phenotype are consistent with those observed in cases of human disease, suggesting that a HFD feeding regime in *Drosophila* larvae can be used to model human disease.

*Drosophila* fed a HFD exhibit many of the same metabolic and physical symptoms as obese humans (Birse et al. 2010; Guida et al. 2019; Hardy et al. 2015). However, existing studies have predominantly focused on transient feeding of adults, and analyzed only females. Our work suggests that larval *Drosophila* provides a better model for chronic dietary treatments, and for the examination of the sex specific effects of diet. We also find that while previous studies have reported most consistent and reproducible effects of HFD treatments on 30% coconut oil supplemented diets (Birse et al. 2010; Diop, Birse, and Bodmer 2017; Guida et al. 2019), there may be variability in tolerance with different genotypes. Our results show that 30% is ideal for inducing cardiac phenotypes in *y^1^w^1118^,* but that a 20% diet is better for *vkgGFP.* This presents an interesting opportunity to examine the effects of genetic background on response to dietary treatment. It has previously been found that a HFD affects heart function due to increased TOR signalling and decreased expression of the lipase Brummer (Birse et al. 2010). These pathways are highly conserved and also altered in human disease. Based on our findings it is possible that different *Drosophila* genotypes have different levels of signalling of these key pathways, therefore altering their ability to respond to the lipotoxic insult of a HFD. Further study can utilize this feeding regime as a model for understanding the genetic basis of differing tolerances to HFD feeding, which may provide insight into how these processes are differentially regulated in human populations.

## Methods

### *Drosophila* strains and dietary treatments

*y^1^w^1118^* (6598, BDSC) and *y^1^w^1118^ ; vkg-GFP (vkg^CC00791^)* lines were used for these experiments (Buszczak et al. 2007) (Bloomington stock centre, NIH P40OD018537). Flies were maintained at room temperature and fed one of 6 different dietary treatments – regular fly food (control), 1.0M sucrose, 10%, 20%, 30%, and 40% coconut oil supplemented (high fat diet). In all dietary treatments the protein source was scaled to match the volume of food. 1.0M sucrose is approximately calorically equivalent to 20% HFD.

All treatments were supplements made to ordinary lab food. Ordinary lab food consists of 3.6L of water, 300g sucrose (0.2M), 150g yeast, 24g KNa tartrate, 3g dipotassium hydrogen orthobasic, 1.5g NaCl, 1.5g CaCl_2,_ 1.5g MgCl_2_, 1.5g ferric sulfur, and 54g of agar. Fly food is autoclaved, cooled to 55°C, then 22mL of 10% tegosept and 15mL of acid mix is added before dispensing. Coconut oil supplements were by volume, sucrose by molarity.

### Triglyceride assay

Triglyceride levels were measured using a serum triglyceride determination kit (Sigma Aldrich, TR0100) (Wat et al. 2020). 5 intact third instar larvae were flash frozen in liquid nitrogen and stored at −80°C before sample preparation. Frozen larvae were ground with a manual homogenizer in 0.1% Tween in PBS. 20µl of buffer per larva was used. Samples were heat treated at 70°C for 10 minutes, then centrifuged at maximum speed for 3 minutes. 10µL of each sample was loaded into a 96 well plate in triplicate. 10µL of a glycerol standard at 2.5mg/mL, 1.25mg/mL, 0.625mg/mL, 0.315mg/mL, 0.156mg/mL, and 0mg/mL were also loaded. 250µL of free glycerol reagent was added to each well, incubated at 37°C, and absorbance was read at 540nm. 50µL of triglyceride reagent was then added, incubated for 10 minutes at 37°C, and absorbance read at 540nm. The change in glycerol levels after addition of the triglyceride reagent was calculated to determine the level of stored triglycerides in the sample. A Bradford assay was then conducted on the same samples and the level of stored triglycerides was divided by the amount of protein in the sample to control for body size.

### Dissections

#### Heart

Dissections were performed by fixing larvae dorsal down to a surface using pins (Brent, Werner, and McCabe 2009). Larvae were bathed in PBS and an incision was made at the ventral midline. The cuticle was pinned back and the gut and fat bodies were removed to reveal the heart. Dissections were performed at third instar, after the onset of wandering behaviour.

#### Fat body

Above process was followed but only the gut was removed to expose the fat bodies.

### Immunohistochemistry

#### Heart

Dissections were fixed for 20 minutes without shaking at room temperature in 4% paraformaldehyde in PBS. Specimens were then washed 3×10 minutes in PBST (0.3% Triton-X-100), before blocking for 30 minutes with NGS (1:15). Primary antibodies were incubated overnight at 4°C with shaking. After incubation with primary 3×10 minute washes in PBST were performed before adding secondary antibodies for one hour at room temperature. Phalloidin was added at the same time as secondary antibodies. Specimens were then washed 3×10 minutes in PBST, with a final wash in PBS to remove detergent. 50% glycerol was added for at least 3 hours, then 70% glycerol overnight. The primary antibody used was mouse anti-Prc (Pericardin, EC11, DSHB, 1:30 dilution). Secondary antibodies used were Alexa Fluor 488 anti-mouse and Alexa Fluor 647 anti-mouse (1:150 dilution). Alexa Fluor 546 and 647 Phalloidin (Thermofisher Scientific) were also used (1:75 dilution).

#### Fat body

Dissections were fixed for 30 minutes at room temperature in 4% paraformaldehyde. Specimens were washed 2×5 minutes in PBST, then incubated in 493/503 BODIPY (1:1000) for 30 minutes. Specimens were then washed 2×5 minutes, placed in 70% glycerol, and immediately mounted for imaging.

### Imaging

A Leica SP5 confocal microscope was used to obtain image stacks. 1µm intervals between frames were used for heart dissections, 0.5µm intervals were used for fat bodies. Fat bodies were imaged from the surface to a depth of 30µm. Hearts were imaged from the ventral face of the cardiac ECM to the dorsal edge of the heart tube. Images were processed using Leica software (LAS AF), ImageJ, and ZEN blue.

### Image quantification and statistics

Pericardin linearity was measured by blind score, with a scale of 1 being a normal meshy matrix, 2 having some linearity but not the majority, and 3 being majority or completely linear.

Pericardin matrix to heart tube ratio was obtained by tracing the outline of the matrix and the heart tube, obtaining areas for each, and generating a ratio.

The percentage of the Collagen-IV matrix that was occupied by clumps was obtained by tracing the total area of the matrix and then tracing the area of each clump within the matrix. The sum of clump areas was then expressed as a percentage of the total matrix area.

Lipid droplet diameter was measured using the line tool in ZEN blue (Zeng et al. 2020).

All measurements from confocal images were performed on unedited images. For publication only images have had brightness and colour balance adjusted using Photoshop CS6.

Statistical analysis of larval health (mass, triglyceride levels, lipid droplet size), Pericardin organization, and diastole/systole were performed using Graphpad Prism (v.9.5.1). Analysis of variance (ANOVA) with a Dunnett’s multiple comparisons test performed. Graphs are plotted with SEM.

For Collagen-IV matrix organization statistics we fit a linear model with the terms sex, diet, and their interaction using the R programming language. Significance of the terms was tested with ANOVA and differences in group means using the emmeans(v1.7.2) package, with 95% confidence intervals reported for all statistics. Sucrose and the equivalent calorie HFD were compared to controls separately to determine if sucrose had a comparable effect to HFD feeding. Because sucrose was not different from controls it was excluded from HFD comparisons. Diastolic and systolic volumes for HFD treatments were fit using a linear model with the term diet to estimate the slope of the line as 95% confidence intervals.

### OCT imaging

Optical coherence tomography (OCT) was used to visualize the heart beating *in vivo* in real time in late third instar larvae. Larvae were adhered to a microscope slide dorsal side up before being placed under the OCT camera. B scans were taken in 3D acquisition mode using a Thorlabs OCT Telesto series TEL221PS system at the widest point of the heart chamber with the following parameters: X size 1257 pixels, 1.03mm, Y size 0, 400 frames, Z field of view 1.2mm. This gives a 20 second video with 20 frames per second. Image stacks were then exported as TIFs and processed in ImageJ (Abràmoff, n.d.). The cross-sectional area was measured at both diastole and systole. Diastolic and systolic volumes across HFD groups were compared using a model estimate with 95% confidence intervals.

## Acknowledgements

We thank João Firmino and the staff of the Centre for Advanced Light Microscopy at McMaster University for technical assistance with OCT.

## Author contributions

R.M.A. – methodology, investigation, visualization, writing; S.N. – investigation; K.P. – statistical analysis, review; J.R.J. – conceptualization, review, editing, supervision, administration, funding.

## Supplemental figures

**Figure S1:**
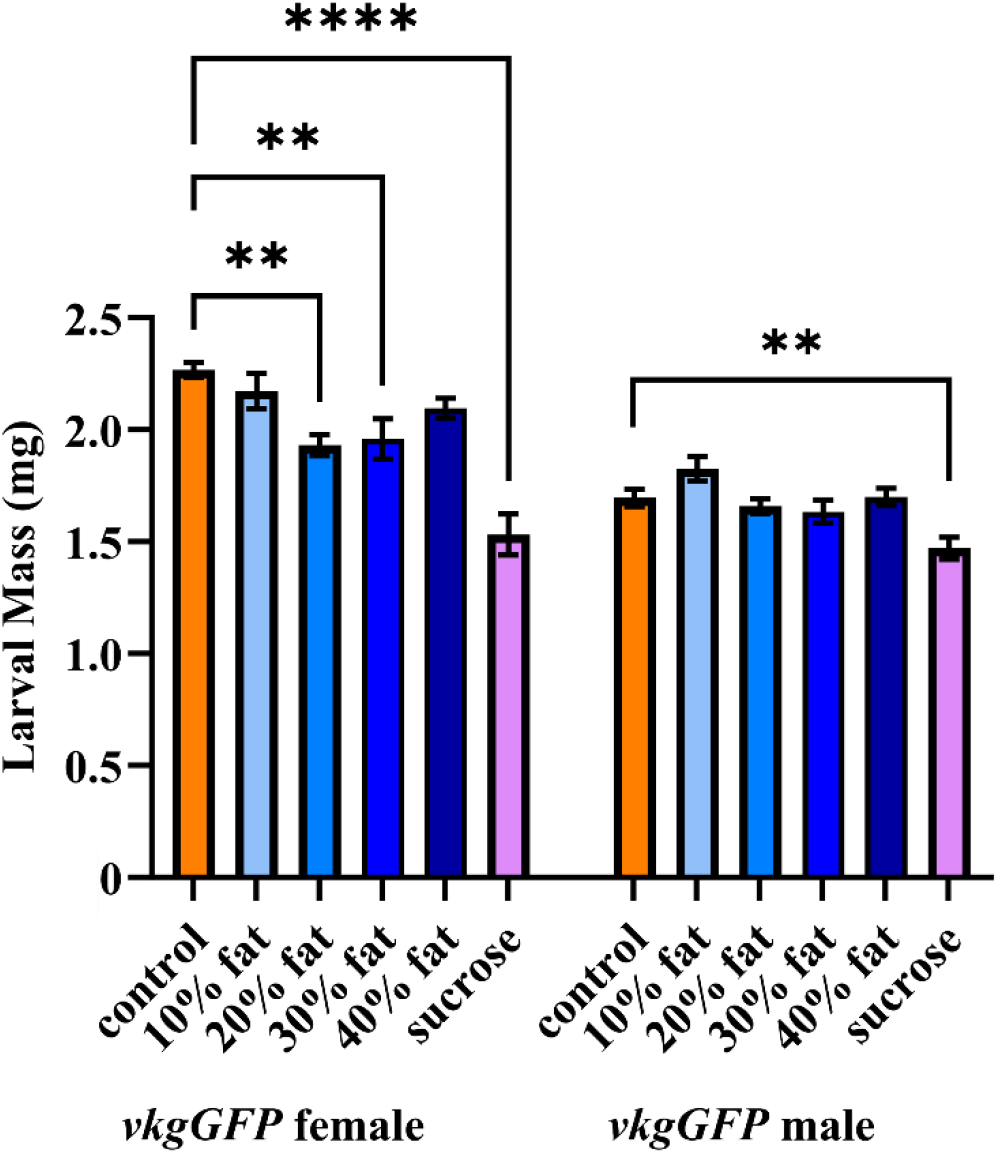
*vkgGFP* larval mass. *vkgGFP* female 20% and 30% high fat diet treatments exhibit lower mass than controls, in males no HFD treatments are significantly reduced compared to controls. The high sucrose diet generates smaller larvae in both males and females. Error bars are SEM. *=p<0.05, **=p<0.01, ****=p<0.0001

**Figure S2:**
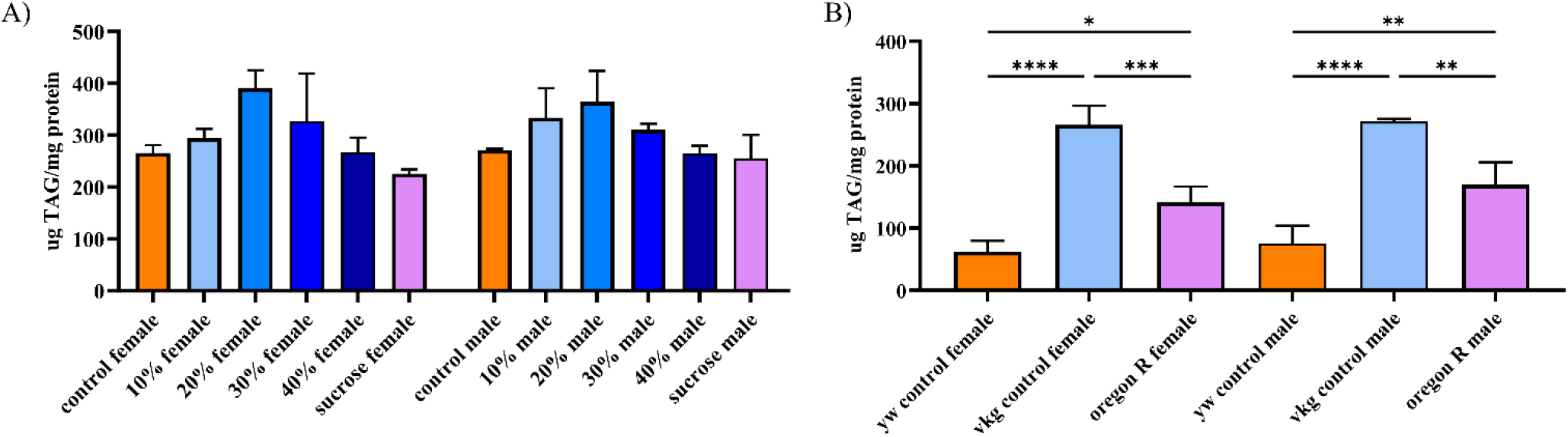
Baseline triglyceride levels vary with genotype. *vkgGFP* larvae demonstrate elevated triglyceride levels at baseline (A). HFD and high sucrose diet treatments do not have an effect on triglyceride levels in this genotype. The baseline level of triglycerides in *y^1^w^1118^, vkgGFP* and Oregon R larvae are variable (B). Both female and male larvae exhibit this variability.

**Figure S3:**
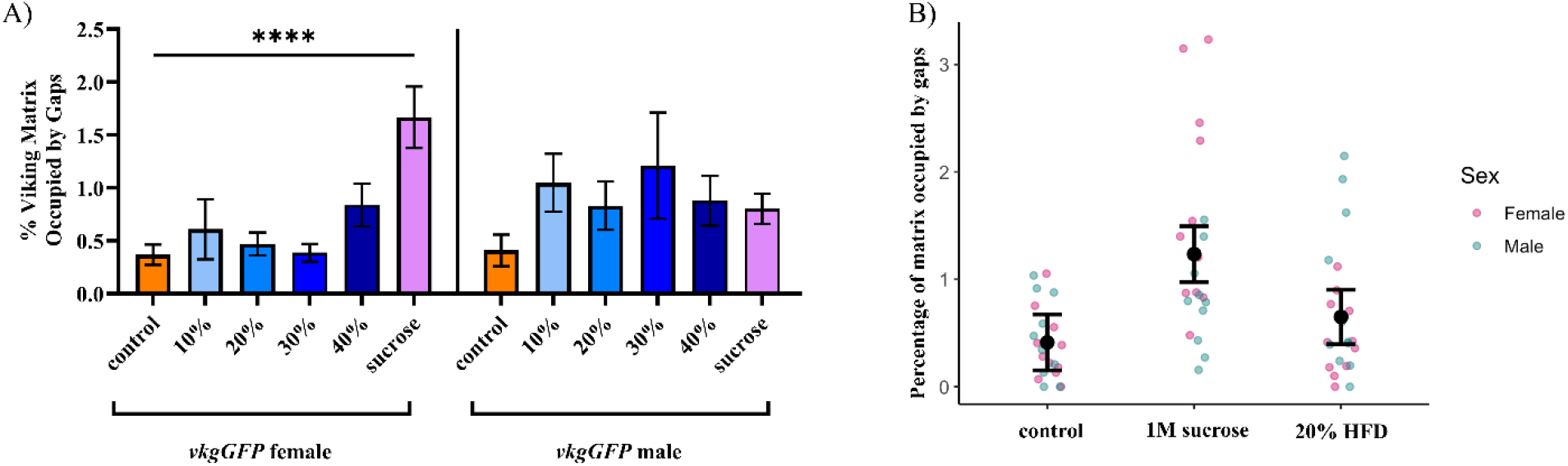
High sucrose diet causes gaps rather than clumps within the *vkgGFP* matrix. The high sucrose diet was found to induce the formation of gaps or holes within the Collagen-IV matrix (A). This phenotype was significant only in females. High fat diets were not found to cause this phenotype (B). Error bars in A are SEM, error bars in B are 95% confidence intervals. ****=p<0.0001

**Figure S4:**
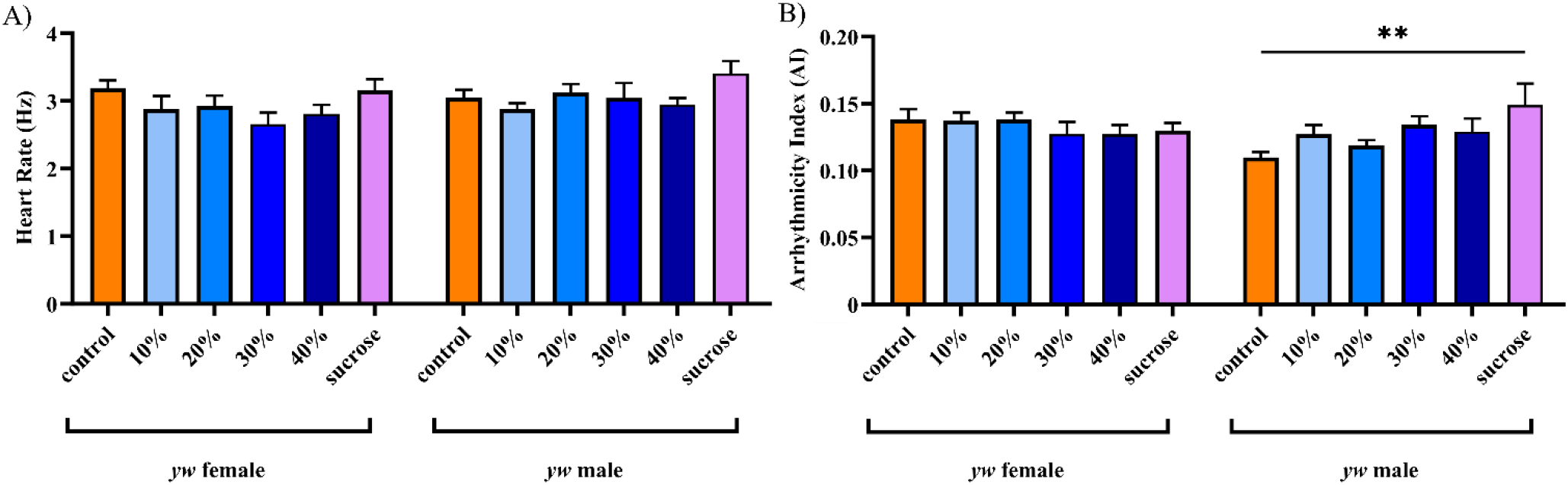
Cardiac functional parameters are unchanged in most dietary treatments. Heart rate was not significantly different with any dietary treatment (A). Arrhythmicity index was unaffected in all treatments except for high sucrose diet males (B). Error bars are SEM. **=p<0.01

## References

1. Abràmoff, Dr Michael D. n.d. “Image Processing with ImageJ.”

2. Alpert, Martin A., Jad Omran, and Brian P. Bostick. 2016. “Effects of Obesity on Cardiovascular Hemodynamics, Cardiac Morphology, and Ventricular Function.” Current Obesity Reports 5 (4): 424–34. https://doi.org/10.1007/s13679-016-0235-6.

3. Berlo, Jop H. van, Onur Kanisicak, Marjorie Maillet, Ronald J. Vagnozzi, Jason Karch, Suh-Chin J. Lin, Ryan C. Middleton, Eduardo Marbán, and Jeffery D. Molkentin. 2014. “C-Kit+ Cells Minimally Contribute Cardiomyocytes to the Heart.” Nature 509 (7500): 337–41. https://doi.org/10.1038/nature13309.

4. Birse, Ryan T., Joan Choi, Kathryn Reardon, Jessica Rodriguez, Suzanne Graham, Soda Diop, Karen Ocorr, Rolf Bodmer, and Sean Oldham. 2010. “High-Fat-Diet-Induced Obesity and Heart Dysfunction Are Regulated by the TOR Pathway in Drosophila.” Cell Metabolism 12 (5): 533–44. https://doi.org/10.1016/j.cmet.2010.09.014.

5. Bonnans, Caroline, Jonathan Chou, and Zena Werb. 2014. “Remodelling the Extracellular Matrix in Development and Disease.” Nature Reviews Molecular Cell Biology 15 (12): 786–801. https://doi.org/10.1038/nrm3904.

6. Brent, Jonathan R., Kristen M. Werner, and Brian D. McCabe. 2009. “Drosophila Larval NMJ Dissection.” Journal of Visualized Experiments : JoVE, no. 24 (February): 1107. https://doi.org/10.3791/1107.

7. Buszczak, Michael, Shelley Paterno, Daniel Lighthouse, Julia Bachman, Jamie Planck, Stephenie Owen, Andrew D Skora, et al. 2007. “The Carnegie Protein Trap Library: A Versatile Tool for Drosophila Developmental Studies.” Genetics 175 (3): 1505–31. https://doi.org/10.1534/genetics.106.065961.

8. Cavalera, Michele, Junhong Wang, and Nikolaos G. Frangogiannis. 2014. “Obesity, Metabolic Dysfunction, and Cardiac Fibrosis: Pathophysiological Pathways, Molecular Mechanisms, and Therapeutic Opportunities.” Translational Research 164 (4): 323–35. https://doi.org/10.1016/j.trsl.2014.05.001.

9. Chartier, Aymeric, Stéphane Zaffran, Martine Astier, Michel Sémériva, and Danielle Gratecos. 2002. “Pericardin, a Drosophila Type IV Collagen-like Protein Is Involved in the Morphogenesis and Maintenance of the Heart Epithelium during Dorsal Ectoderm Closure.” Development 129 (13): 3241–53. https://doi.org/10.1242/dev.129.13.3241.

10. Cox, Thomas R., and Janine T. Erler. 2011. “Remodeling and Homeostasis of the Extracellular Matrix: Implications for Fibrotic Diseases and Cancer.” Disease Models & Mechanisms 4 (2): 165–78. https://doi.org/10.1242/dmm.004077.

11. Diop, Soda Balla, Ryan T. Birse, and Rolf Bodmer. 2017. “High Fat Diet Feeding and High Throughput Triacylglyceride Assay in Drosophila Melanogaster.” Journal of Visualized Experiments : JoVE, no. 127 (September): 56029. https://doi.org/10.3791/56029.

12. Drechsler, Maik, Ariane C. Schmidt, Heiko Meyer, and Achim Paululat. 2013. “The Conserved ADAMTS-like Protein Lonely Heart Mediates Matrix Formation and Cardiac Tissue Integrity.” Edited by Norbert Perrimon. PLoS Genetics 9 (7): e1003616. https://doi.org/10.1371/journal.pgen.1003616.

13. El Hajj, Elia C., Milad C. El Hajj, Van K. Ninh, and Jason D. Gardner. 2018. “Inhibitor of Lysyl Oxidase Improves Cardiac Function and the Collagen/MMP Profile in Response to Volume Overload.” American Journal of Physiology. Heart and Circulatory Physiology 315 (3): H463–73. https://doi.org/10.1152/ajpheart.00086.2018.

14. Fidler, Aaron L, Carl E Darris, Sergei V Chetyrkin, Vadim K Pedchenko, Sergei P Boudko, Kyle L Brown, W Gray Jerome, Julie K Hudson, Antonis Rokas, and Billy G Hudson. 2017. “Collagen IV and Basement Membrane at the Evolutionary Dawn of Metazoan Tissues.” Edited by Harry C Dietz. ELife 6 (April): e24176. https://doi.org/10.7554/eLife.24176.

15. Garrett, Sara M., Eileen Hsu, Justin M. Thomas, Joseph M. Pilewski, and Carol Feghali-Bostwick. 2019. “Insulin-like Growth Factor (IGF)-II-Mediated Fibrosis in Pathogenic Lung Conditions.” PLOS ONE 14 (11): e0225422. https://doi.org/10.1371/journal.pone.0225422.

16. Guida, Maria Clara, Ryan Tyge Birse, Alessandra Dall’Agnese, Paula Coutinho Toto, Soda Balla Diop, Antonello Mai, Peter D. Adams, Pier Lorenzo Puri, and Rolf Bodmer. 2019. “Intergenerational Inheritance of High Fat Diet-Induced Cardiac Lipotoxicity in Drosophila.” Nature Communications 10 (January): 193. https://doi.org/10.1038/s41467-018-08128-3.

17. Hardy, Christopher M., Ryan T. Birse, Matthew J. Wolf, Lin Yu, Rolf Bodmer, and Allen G. Gibbs. 2015. “Obesity-Associated Cardiac Dysfunction in Starvation-Selected Drosophila Melanogaster.” American Journal of Physiology-Regulatory, Integrative and Comparative Physiology 309 (6): R658–67. https://doi.org/10.1152/ajpregu.00160.2015.

18. Hartley, Paul S., Khatereh Motamedchaboki, Rolf Bodmer, and Karen Ocorr. 2016. “SPARC– Dependent Cardiomyopathy in Drosophila.” Circulation. Cardiovascular Genetics 9 (2): 119–29. https://doi.org/10.1161/CIRCGENETICS.115.001254.

19. Hinderer, Svenja, and Katja Schenke-Layland. 2019. “Cardiac Fibrosis – A Short Review of Causes and Therapeutic Strategies.” Advanced Drug Delivery Reviews 146 (June): 77–82. https://doi.org/10.1016/j.addr.2019.05.011.

20. Howard, Angela M., Kimberly S. LaFever, Aidan M. Fenix, Cherie’ R. Scurrah, Ken S. Lau, Dylan T. Burnette, Gautam Bhave, Nicholas Ferrell, and Andrea Page-McCaw. 2019. “DSS-Induced Damage to Basement Membranes Is Repaired by Matrix Replacement and Crosslinking.” Journal of Cell Science, January, jcs.226860. https://doi.org/10.1242/jcs.226860.

21. Hughes, C. J. R., and J. Roger Jacobs. 2017. “Dissecting the Role of the Extracellular Matrix in Heart Disease: Lessons from the Drosophila Genetic Model.” Veterinary Sciences 4 (2): 24. https://doi.org/10.3390/vetsci4020024.

22. Hughes, C.J.R., S. Turner, R.M. Andrews, A. Vitkin, and J.R. Jacobs. 2020. “Matrix Metalloproteinases Regulate ECM Accumulation but Not Larval Heart Growth in Drosophila Melanogaster.” Journal of Molecular and Cellular Cardiology 140 (March): 42–55. https://doi.org/10.1016/j.yjmcc.2020.02.008.

23. Jourdan-LeSaux, Claude, Jianhua Zhang, and Merry L. Lindsey. 2010. “Extracellular Matrix Roles during Cardiac Repair.” Life Sciences 87 (13): 391–400. https://doi.org/10.1016/j.lfs.2010.07.010.

24. Meschiari, Cesar A., Osasere Kelvin Ero, Haihui Pan, Toren Finkel, and Merry L. Lindsey. 2017. “The Impact of Aging on Cardiac Extracellular Matrix.” GeroScience 39 (1): 7–18. https://doi.org/10.1007/s11357-017-9959-9.

25. Mouw, Janna K., Guanqing Ou, and Valerie M. Weaver. 2014. “Extracellular Matrix Assembly: A Multiscale Deconstruction.” Nature Reviews Molecular Cell Biology 15 (12): 771–85. https://doi.org/10.1038/nrm3902.

26. Na, Jianbo, Laura Palanker Musselman, Jay Pendse, Thomas J. Baranski, Rolf Bodmer, Karen Ocorr, and Ross Cagan. 2013. “A Drosophila Model of High Sugar Diet-Induced Cardiomyopathy.” Edited by Eric Rulifson. PLoS Genetics 9 (1): e1003175. https://doi.org/10.1371/journal.pgen.1003175.

27. Nishida, Kazuhiko, and Kinya Otsu. 2017. “Inflammation and Metabolic Cardiomyopathy.” Cardiovascular Research 113 (4): 389–98. https://doi.org/10.1093/cvr/cvx012.

28. Palanker Musselman, Laura, Jill L. Fink, Kirk Narzinski, Prasanna Venkatesh Ramachandran, Sumitha Sukumar Hathiramani, Ross L. Cagan, and Thomas J. Baranski. 2011. “A High-Sugar Diet Produces Obesity and Insulin Resistance in Wild-Type *Drosophila*.” Disease Models & Mechanisms 4 (6): 842–49. https://doi.org/10.1242/dmm.007948.

29. Pastor-Pareja, José C. 2020. “Atypical Basement Membranes and Basement Membrane Diversity – What Is Normal Anyway?” Journal of Cell Science 133 (8): jcs241794. https://doi.org/10.1242/jcs.241794.

30. Pehrsson, Martin, Joachim Høg Mortensen, Tina Manon-Jensen, Anne-Christine Bay-Jensen, Morten Asser Karsdal, and Michael Jonathan Davies. 2021. “Enzymatic Cross-Linking of Collagens in Organ Fibrosis – Resolution and Assessment.” Expert Review of Molecular Diagnostics 21 (10): 1049–64. https://doi.org/10.1080/14737159.2021.1962711.

31. Rewitz, Kim F., Naoki Yamanaka, and Michael B. O’Connor. 2013. “Chapter One - Developmental Checkpoints and Feedback Circuits Time Insect Maturation.” In Current Topics in Developmental Biology, edited by Yun-Bo Shi, 103:1–33. Animal Metamorphosis. Academic Press. https://doi.org/10.1016/B978-0-12-385979-2.00001-0.

32. Rotstein, Barbara, and Achim Paululat. 2016. “On the Morphology of the Drosophila Heart.” Journal of Cardiovascular Development and Disease 3 (2): 15. https://doi.org/10.3390/jcdd3020015.

33. Sessions, Ayla O., Gaurav Kaushik, Sarah Parker, Koen Raedschelders, Rolf Bodmer, Jennifer E. Van Eyk, and Adam J. Engler. 2017. “Extracellular Matrix Downregulation in the Drosophila Heart Preserves Contractile Function and Improves Lifespan.” Matrix Biology 62 (October): 15–27. https://doi.org/10.1016/j.matbio.2016.10.008.

34. Sidney, Stephen, Catherine Lee, Jennifer Liu, Sadiya S. Khan, Donald M. Lloyd-Jones, and Jamal S. Rana. 2022. “Age-Adjusted Mortality Rates and Age and Risk–Associated Contributions to Change in Heart Disease and Stroke Mortality, 2011-2019 and 2019-2020.” JAMA Network Open 5 (3): e223872. https://doi.org/10.1001/jamanetworkopen.2022.3872.

35. Tian, Gang, Chaodi Luo, and Lei Liu. 2022. “Epicardial Adipose Tissie-Derived Leptin Induce Mmps/Timps Imbalance and Promote Cardiac Fibrosis through Jak2/Ros/Na/k-Atpase/Erk1/2 Signaling Pathway in High Fat Diet-Induced Obese Rats.” Journal of the American College of Cardiology 79 (9_Supplement): 1544–1544. https://doi.org/10.1016/S0735-1097(22)02535-9.

36. Travers, Joshua G., Fadia A. Kamal, Jeffrey Robbins, Katherine E. Yutzey, and Burns C. Blaxall. 2016. “Cardiac Fibrosis: The Fibroblast Awakens.” Circulation Research 118 (6): 1021–40. https://doi.org/10.1161/CIRCRESAHA.115.306565.

37. Vujic, Ana, Niranjana Natarajan, and Richard T. Lee. 2020. “Molecular Mechanisms of Heart Regeneration.” Seminars in Cell & Developmental Biology 100 (April): 20–28. https://doi.org/10.1016/j.semcdb.2019.09.003.

38. Walker, C. A., and F. G. Spinale. 1999. “The Structure and Function of the Cardiac Myocyte: A Review of Fundamental Concepts.” The Journal of Thoracic and Cardiovascular Surgery 118 (2): 375–82. https://doi.org/10.1016/S0022-5223(99)70233-3.

39. Walls, Stanley M., Dale A. Chatfield, Karen Ocorr, Greg L. Harris, and Rolf Bodmer. 2020. “Systemic and Heart Autonomous Effects of Sphingosine Δ-4 Desaturase Deficiency in Lipotoxic Cardiac Pathophysiology.” Disease Models & Mechanisms, January, dmm.043083. https://doi.org/10.1242/dmm.043083.

40. Wat, Lianna W., Charlotte Chao, Rachael Bartlett, Justin L. Buchanan, Jason W. Millington, Hui Ju Chih, Zahid S. Chowdhury, et al. 2020. “A Role for Triglyceride Lipase Brummer in the Regulation of Sex Differences in Drosophila Fat Storage and Breakdown.” Edited by Bassem A. Hassan. PLOS Biology 18 (1): e3000595. https://doi.org/10.1371/journal.pbio.3000595.

41. Zeng, Jie, Nhan Huynh, Brian Phelps, and Kirst King-Jones. 2020. “Snail Synchronizes Endocycling in a TOR-Dependent Manner to Coordinate Entry and Escape from Endoreplication Pausing during the Drosophila Critical Weight Checkpoint.” Edited by Bruce Edgar. PLOS Biology 18 (2): e3000609. https://doi.org/10.1371/journal.pbio.3000609.

